# Retrospective analysis of the use of osteoporosis medication at the presentation of fragility fractures in a predominantly Hispanic population

**DOI:** 10.1101/2020.12.24.424289

**Authors:** Michael J. Serra-Torres, Annelyn Torres-Reveron

**Author notes:** Corresponding Author: A. Torres-Reveron, PhD.

## Abstract

Despite the high incidence of osteoporosis, many patients at risk of fragility fractures may not initiate treatment due to concerns about side effects, or cost. We aimed to identify whether the patient population presenting with fragility fractures is receiving prophylactic treatment for osteoporosis within a single academic hospital in the southernmost region of the United States. This region is characterized by a high number of patients from Hispanic/Latino heritage (80%) and reduced access to healthcare services. We conducted a three-year, retrospective cohort study of patients presenting with low impact fractures of the humerus or the shoulder griddle, lower end of radius or ulna and forearm, hip fractures (femoral neck, intertrochanteric/subtrochanteric), the lower end of tibia, medial malleolus, lateral malleolus or lower leg. Male and female subjects of 50 years or older were included. Demographic data and information on medications reported at fracture presentation were extracted from electronic medical records. We found that 42% of the patients were taking at least one medication to prevent osteoporosis. The predominant combination was vitamin D plus calcium and bisphosphonates. If patients taking only vitamin D plus calcium are excluded, 16.7% of the sample took osteoporosis medications at the fragility fracture presentation. The likelihood of taking osteoporosis medication was increased by age and type of health insurance (Medicare/private insurance), and concomitant diagnosis of impaired gait and mobility. The percentage of the patients taking prophylactic medications for osteoporosis at the time of a fragility fracture was comparable to reported national standards and associated with increased age and health insurance coverage. In a predominantly Hispanic/Latino patient population living in a medically underserved region, there is substantial recognition and prevention strategies for osteoporosis.

## Introduction

Osteoporosis is a systemic bone disease characterized by a decrease in bone strength reflected by a combination of bone mineral density and bone quality, which includes architecture, turnover, damage accumulation, matrix mineralization, and collagen composition [1]. It affects one in four women and one in 8 men over the age of 50 years. It is estimated that 44 million people have either osteoporosis or low bone mass in the United States, representing 55% of the population 50 years and older [2]. The annual cost of osteoporosis fractures in 2018 is estimated at $57 billion dollars and is projected to reach $95 billion by the year 2040 [3–5]. Osteoporosis can be primarily influenced by lifestyle changes and the use of prophylactic bone-forming/preserving agents. Examining the frequency and use of prophylactic medications to prevent osteoporotic/fragility fractures can help design interventions to reduce negative impacts on the elderly.

Osteoporosis leads to fragility fractures most commonly observed in the distal radius, proximal humerus, vertebral, proximal femur, and ankle that result from minimal trauma, such as a fall from a standing height. Fragility fractures can lead to loss of functionality, pain, limitation of activities of daily living (ADLs), increased morbidity, disability, and significant adverse effect on the quality of life [6]. These consequences present a high burden on the healthcare systems due to increased hospitalizations, surgeries, utilization requirements (e.g., Rehabilitation/P.T., home care, nursing), and medical costs [3]. Fragility fractures account for more than 40% of the hospital admissions of women 55 years or over. In the United States, the annual cost for fragility fracture admissions was more significant than the cost for myocardial infarctions, stroke, and breast cancer [7]. It is estimated that the global number of fragility fractures was 9 million annually in 2000, including 1.6 million hip fractures, 1.7 million wrist fractures, and 1.4 million spine fractures [2]. It is predicted that this number will continue to increase due to the changes in life expectancy and to change demographics globally. Moreover, by the year 2050, it is estimated that 70% of all patients with hip fractures will be located in Asia, Latin America, and the Middle East [8].

In 2020, the United States population of patients over the age of 50 years old is between 13-20%. This number is projected to increase to 28-40% by 2050. Hispanics, according to the 2000s United States-Census, represent the largest minority group at 12.5% of the total population, and this ethnic group is anticipated to increase to 25% by the year 2050 (12). Concurrently, the incidence of hip fractures in Latin Americans is projected to increase to 12.5% by 2050, representing an incidence increase of 5.5% from that in 1990 [9]. Despite these projections, there is a lack of research and data assessing the risk factors for osteoporosis and osteopenia in Hispanics [10–12]. The risk factors for osteoporosis can be classified as modifiable, such as smoking, low body weight, hormone deficiency, alcohol intake, physical activity, dietary intake, falls, medications, chronic conditions, social clinical and medical insurance status, and non-modifiable risk factors: race, age, sex, dementia, past medical and family history of fractures [13]. Understanding the interaction of modifiable and non-modifiable factors in Hispanics is crucial to design and implement appropriate prevention programs for osteoporosis.

Osteoporosis continues to be under-recognized and under-treated despite its massive economic cost and impact on morbidity and mortality [14]. The viability of health systems payer (insurance) and economic conditions has been related to quality in health care provided in Latin American regions[3]. A recent study showed that in Latin America, 43.7% of postmenopausal women had not received osteoporosis therapy, whereas 14.3% of women in the United States and 11.3% in Europe had received therapy [15,16].

Most medications for treatment of osteoporosis work by either decreasing bone resorption, (bisphosphonates, selective estrogen receptor modulators, Denosumab-RANK ligand inhibitors) or increasing bone formation (recombinant parathyroid hormone) [11,12]. Using a patient cohort from the southernmost border of the United States composed of approximately 80% Hispanics, we aim to recognize whether the population presenting with fragility fractures is receiving prophylactic treatment for such condition. Based on prior reports from Latin America, we hypothesized that using prophylactic medications for osteoporosis is significantly lower in our cohort than previously reported national studies in the United States. Our target is to improve the knowledge of osteoporosis treatment use in the Hispanic population at presentation with a fragility fracture and increase early interventions in communities at risk.

## Methods

### Study Design and Setting

This is a three-year, retrospective cohort study of patients from a single hospital, Level 2 Trauma Center in South Texas. The study conformed to the Declaration of Helsinki and the U.S. Federal Policy for the Protection of Humans Subjects and was approved by the Institutional Review Board. The study team submitted a full waiver of authorization under the Health Insurance Portability and Accountability Act (HIPAA, 1996) and was approved by the Institutional Review Board to conduct this retrospective study. The retrospective period of chart review was set from January 1, 2017, to December 31, 2019.

### Subjects

Male and female subjects 50 years and older presenting to any of the clinics or emergency rooms at our health system were included in the study. The following types of fractures were included: upper end of the humerus or the shoulder griddle, lower end of radius or ulna and forearm, displaced or non-displaced femoral neck, intertrochanteric/subtrochanteric fractures, the lower end of tibia, medial malleolus, lateral malleolus, or lower leg. Patients were excluded from the study if the onset of the injury or presentation was outside the study period, if the fracture was already treated at another institution (e.g., referred to a rehabilitation hospital), or if the fracture was nonunion. Additional exclusion criteria were based on the mechanism of injury: fractures produced as a result of projectiles, motor vehicle accidents, or from a height higher than 1 meter were excluded from the cohort. Fractures due to diagnosed/documented cancer were excluded as well. Fractures of the vertebrae were purposefully excluded due to the documented underdiagnoses [17,18] of these types of injuries, thus potentially creating a selection bias.

### Variables

The demographic variables included were age, sex, ethnicity, body mass index, date of arrival, presentation site, and health insurance or self-pay. From the medical record, we obtained the onset of the injury, the primary diagnosis and secondary diagnosis, the mechanism of injury, treatment received and date, intensive care unit (ICU) usage, comorbidities, home medications use at the time of presentation, zip code of primary residency, discharge disposition, and history of prior dual-energy X-ray absorptiometry (DXA) scan. The type and number of drugs for osteoporosis treatment and prevention were classified based on their mechanism of action: biphosphonates, RANK ligand inhibitors, estrogen, selective estrogen receptor modulators, sclerostin inhibitor, parathyroid hormone analog, calcium, and vitamin D. As a surrogate for patient’s mortality at six months and 12 months, we used the last known reported activity at the hospital, regardless of services requested. In some cases, detailed notes from clinicians indicated the patient’s status.

### Data source/measurements

Patients meeting inclusion criteria were identified via an electronic report from the hospital trauma databank and the business intelligence department at our hospital system. Once the medical record numbers were obtained, the same variables were extracted from all patient’s records. All presented information was part of the subject’s standard of care as documented in their medical record, and no data were collected directly from the patient. To maintain quality and consistency in the data extraction process, a data analyst filtered the information about home medications, and researchers verified the results. An orthopedic surgeon provided oversight of the data collection process and designed the group comparisons.

## Statistical Methods

Descriptive statistics were used for the entire study population. Frequencies and column percentages were used to summarize categorical variables. The normal distribution of continuous variables was measured using the Shapiro-Wilk goodness-of-fit test. Non-normally distributed variables were analyzed using the Wilcoxon test, and normally distributed variables were analyzed using the Student t-test for independent samples. Chi-square or Fisher exact tests were used for categorical variables. Multinomial regression analyses were used to explore the changes in medication across injury types, taking into consideration the age and sex of the patients. The statistical analyses were two-sided and conducted using JMP 15.0 (SAS Institute, Inc, Carry, NC, USA). The statistical significance was set at p< 0.05 and the power was calculated at

## Results

### Participants

A total of 864 cases of fractures in patients older than 50 years of age were initially identified: 121 shoulder cases, 230 wrist cases, 297 hip cases, and 216 ankle cases. Most of the excluded clinical cases were due to the mechanisms of high energy injuries: pedestrian-MVA injuries, motor vehicle accidents, and falls from a height greater than 1 meter. A handful of cases were excluded due to inaccurate coding of fracture. The final cohort consisted of 719 patients.

### Demographic characteristics and medication use

Table 1 describes the number of cases included in each cohort grouped by the site of injury. Patients were predominantly female (>60%), except for shoulder injuries, where 16% of the cases were females. Eighty-one percent of the patients self-reported as Hispanic/Latino ethnicity, which is representative of the demographic distribution for the region. The average age of the cohort was 74.03 ± 11.90 years. Patients with ankle injuries were the youngest in the cohort, with an average age of 66.72 ± 11.08. The length of stay in the hospital was highest for hip injuries (3.19 ± 5.05 days) and lowest for wrist injuries (0.32 ± 1.87 days). Only 62 patients (8.6%) had a prior diagnosis of osteoporosis. The highest percentage of prior diagnosis was in patients with hip fractures and the lowest was in patients with ankle fractures (13.1% and 5.3%, respectively). Of the patients with prior diagnosis, 51 (82%) were taking osteoporosis medications, while 11 (17.7%) were not taking any osteoporosis medication. 98% of the patients diagnosed with osteoporosis were female, while only one patient was male. 10% of the cohort (72 patients) had a dual-energy X-ray absorptiometry (DXA) scan, from which 66 patients were female, and only 6 were male. The highest percentage of patients who received a DXA presented with shoulder fractures (12%).

**Table 1:** Demographics and baseline characteristics.

| Variable | All | Wrist | Shoulder | Hip | Ankle |
| --- | --- | --- | --- | --- | --- |
| N. of patients | 719 | 197 | 99 | 236 | 187 |
| Gender Female, n (%) | 550 (76.5) | 160 (81.2) | 16 (16.2) | 165 (69.9) | 142 (75.9) |
| Hispanic/Latino Ethnicity, n (%) | 588 (81.8) | 174 (88.3) | 80 (80.8) | 181 (76.7) | 153 (81.8) |
| Age, mean (SD) | 74.03 (11.90) | 72.07 (11.41) | 75.6 (9.42) | 80.80 (9.83) | 66.72 (11.08) |
| LOS, mean (SD) | 1.38 (3.57) | 0.32 (1.87) | 1.06 (2.49) | 3.19 (5.05) | 0.37 (1.87) |
| Prior osteoporosis Dx, n (% of total) | 62 (8.6) | 13 (6.6) | 8 (8.0) | 31 (13.1) | 10 (5.3) |
| Prior DXA, n (% of total) | 72 (10.0) | 21 (10.7) | 12 (12.1) | 20 (8.5) | 19 (10.2) |

N, number; Dx, diagnosis; SD, standard deviation; DXA, dual energy X-ray absorptiometry

Patients were stratified based on the number of medications for osteoporosis they were taking (0 to 4). Forty-two percent of the patients, regardless of the fracture site, were taking at least one medication, while 58% were not taking any medication for osteoporosis at the time of fracture (Table 2). Within those taking medications, the vast majority took one or two. The most common medications were a combination of Vitamin D plus calcium and bisphosphonates (Table 3). Other than Vitamin-D plus calcium, only 16.7% were on medications for osteoporosis. The percentage of patients taking medications also varied by fracture site, with the highest percentage of patients in the hip fracture group (58.9%) and the lowest in the ankle fractures group (27.8%). Patients who had Medicare as the principal payer constituted 70% of the cohort, which is expected due to the age group. The type of health insurance influenced whether the patients took osteoporosis medication (X^2^= 66.78, d.f.= 4, p< 0.001; Fig. 1). The odds of taking osteoporosis medication for patients on Medicare compared to self-pay was 6.84 (95% CI: 2.62 to 17.85). The odds were even higher for patients in Medicare than other payment types grouped together (government, charity, indigent): 11.76 (95% CI: 3.56 to 38.86). However, the odds of taking osteoporosis medications for those patients in Medicare compared to private insurance was similar at 1.49 (95% CI: 0.55 to 4.08; Fig. 1).

**Table 2:** Number and type of medications reported at fragility fracture presentation.

| Variable | All | Wrist | Shoulder | Hip | Ankle |
| --- | --- | --- | --- | --- | --- |
| **Number of osteoporosis medications reported at presentation, n (% of column)** |  |  |  |  |  |
| None | 411 (57.1) | 125 (63.4) | 54 (54.5) | 97 (41.1) | 135 (72.1) |
| 1 | 157 (21.8) | 42 (21.3) | 29 (29.3) | 56 (23.7) | 30 (16.0) |
| 2 | 111 (15.4) | 21 (10.7) | 13 (13.1) | 59 (25.0) | 18 (9.6) |
| 3 | 34 (4.7) | 7 (3.5) | 2 (2.0) | 21 (8.9) | 4 (2.1) |
| 4 | 6 (0.8) | 2 (1.0) | 1 (1.0) | 3 (1.3) | 0 |
| **Number of osteoporosis medications reported at presentation, *excluding calcium and vitamin D* n (% of column)** |  |  |  |  |  |
| None | 599 (83.3) | 164 (83.3) | 85 (85.9) | 190 (85.5) | 160 (85.6) |
| 1 | 107 (14.9) | 31 (15.7) | 10 (10.1) | 42 (17.8) | 24 (12.8) |
| 2 | 12 (1.7) | 2 (1.0) | 4 (4.0) | 3 (1.3) | 3 (1.6) |
| 3 | 1 (0.14) | 0 | 0 | 1 (0.4) | 0 |

| Medication class reported at presentation, *excluding calcium and vitamin D*, n (% of column) |  |  |  |  |  |
| --- | --- | --- | --- | --- | --- |
| Biphosphonates | 90 (68.2) | 24 (61.5) | 12 (9.1) | 40 (30.3) | 14 (10.6) |
| Estrogen | 19 (14.4) | 8 (20.5) | 2 (11.1) | 3 (5.8) | 6 (26.1) |
| Denosumab | 11 (8.3) | 4 (10.3) | 1 (5.6) | 5 (9.6) | 1 (4.3) |
| Raloxifene | 6 (4.5) | 2 (5.1) | 2 (11.2) | 1 (1.9) | 1 (4.3) |
| Calcitonin | 6 (4.5) | 1 (2.6) | 1 (5.6) | 3 (5.8) | 1 (4.3) |

**Table 3:**
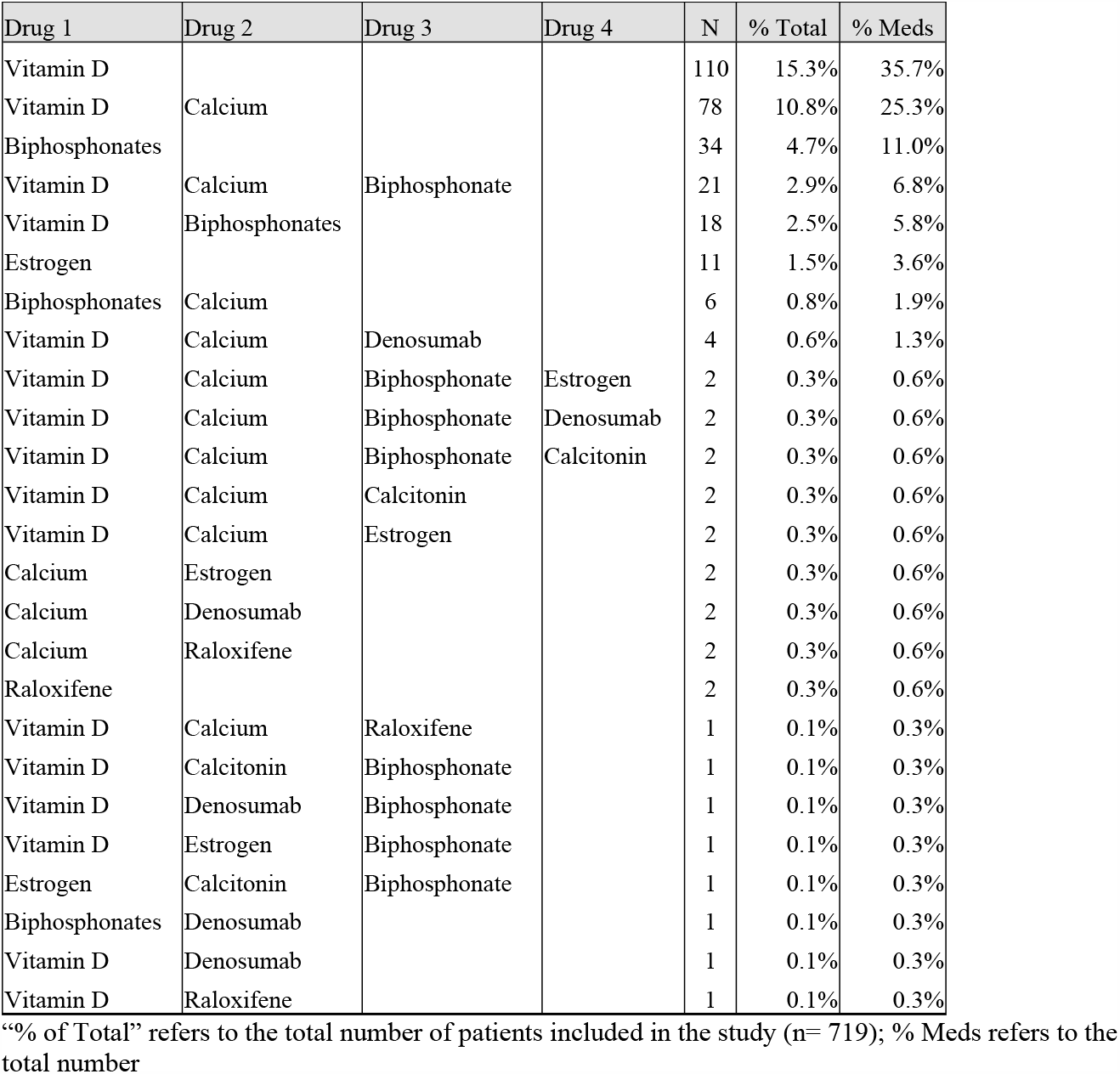
Combinations of osteoporosis medications used from the most frequent to the least frequent.

| Drug 1 | Drug 2 | Drug 3 | Drug 4 | N | % Total | % Meds |
| --- | --- | --- | --- | --- | --- | --- |
| Vitamin D |  |  |  | 110 | 15.3% | 35.7% |
| Vitamin D | Calcium |  |  | 78 | 10.8% | 25.3% |
| Biphosphonates |  |  |  | 34 | 4.7% | 11.0% |
| Vitamin D | Calcium | Biphosphonate |  | 21 | 2.9% | 6.8% |
| Vitamin D | Biphosphonates |  |  | 18 | 2.5% | 5.8% |
| Estrogen |  |  |  | 11 | 1.5% | 3.6% |
| Biphosphonates | Calcium |  |  | 6 | 0.8% | 1.9% |
| Vitamin D | Calcium | Denosumab |  | 4 | 0.6% | 1.3% |
| Vitamin D | Calcium | Biphosphonate | Estrogen | 2 | 0.3% | 0.6% |
| Vitamin D | Calcium | Biphosphonate | Denosumab | 2 | 0.3% | 0.6% |
| Vitamin D | Calcium | Biphosphonate | Calcitonin | 2 | 0.3% | 0.6% |
| Vitamin D | Calcium | Calcitonin |  | 2 | 0.3% | 0.6% |
| Vitamin D | Calcium | Estrogen |  | 2 | 0.3% | 0.6% |
| Calcium | Estrogen |  |  | 2 | 0.3% | 0.6% |
| Calcium | Denosumab |  |  | 2 | 0.3% | 0.6% |
| Calcium | Raloxifene |  |  | 2 | 0.3% | 0.6% |
| Raloxifene |  |  |  | 2 | 0.3% | 0.6% |
| Vitamin D | Calcium | Raloxifene |  | 1 | 0.1% | 0.3% |
| Vitamin D | Calcitonin | Biphosphonate |  | 1 | 0.1% | 0.3% |
| Vitamin D | Denosumab | Biphosphonate |  | 1 | 0.1% | 0.3% |
| Vitamin D | Estrogen | Biphosphonate |  | 1 | 0.1% | 0.3% |
| Estrogen | Calcitonin | Biphosphonate |  | 1 | 0.1% | 0.3% |
| Biphosphonates | Denosumab |  |  | 1 | 0.1% | 0.3% |
| Vitamin D | Denosumab |  |  | 1 | 0.1% | 0.3% |
| Vitamin D | Raloxifene |  |  | 1 | 0.1% | 0.3% |

“% of Total” refers to the total number of patients included in the study (n= 719); % Meds refers to the total number

**Fig. 1:**
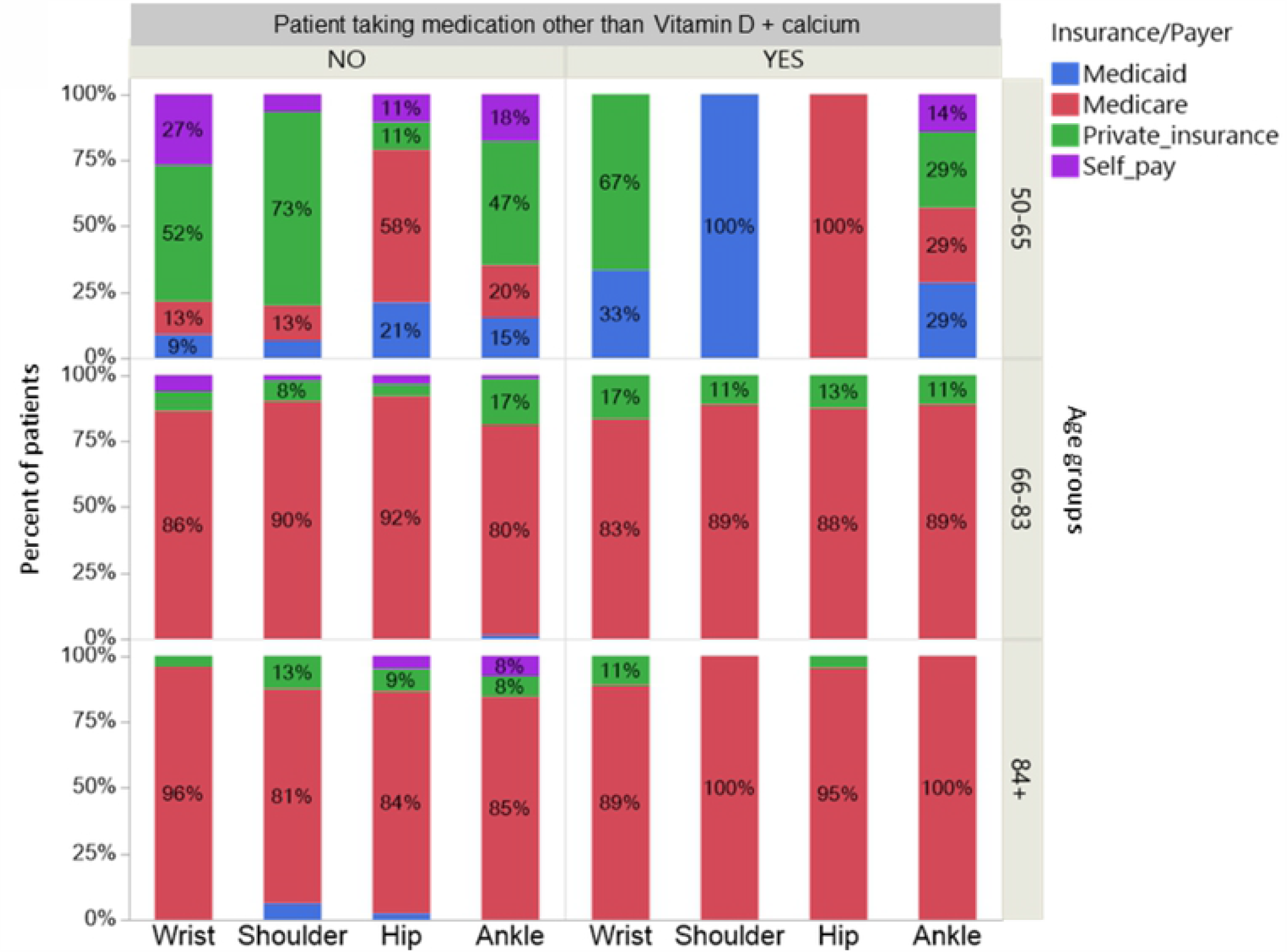
Percent of patients taking osteoporosis medications, with vitamin D plus calcium excluded, by the type of health insurance or payer. Patients who took medications for osteoporosis (YES column) had predominantly Medicare or Medicaid. However, at ages younger than 66, many patients were not taking any medications for osteoporosis (NO column) regardless of the high incidence of private insurance.

Besides a documented diagnosis for osteoporosis, ten other comorbidities were collected from the patient charts: hypertension, obesity, impaired gait or mobility, diabetes, hyperlipidemia, osteoarthritis, other cardiovascular conditions besides hypertension, thyroid-related disease, renal disease, and any cancer. The three most frequent comorbidities in the cohort were: hypertension (59.1%), obesity (47.7%), and impaired gait or mobility (45.2%).

Neither hypertension nor obesity influenced whether the patient was taking any type of osteoporosis medications. However, patients with impaired gait or mobility and patients with thyroid-related disease had an increased odds ratio of receiving osteoporosis medications at 2.56 (95% CI, 1.80 to 3.65) and 5.33 (95% CI, 3.04 to 9.36), respectively. The remaining comorbidities did not influence osteoporosis medication patterns.

### Outcomes-based on sex and age group

To further understand osteoporosis medication use patterns across sex, data were stratified based on three age groups: 50-65, 66-83, and 83 years and older (Fig. 2). The age groups were constructed using the 25^th^ quartile (65 years) and 75^th^ quartile (83 years) for age.

**Fig. 2:**
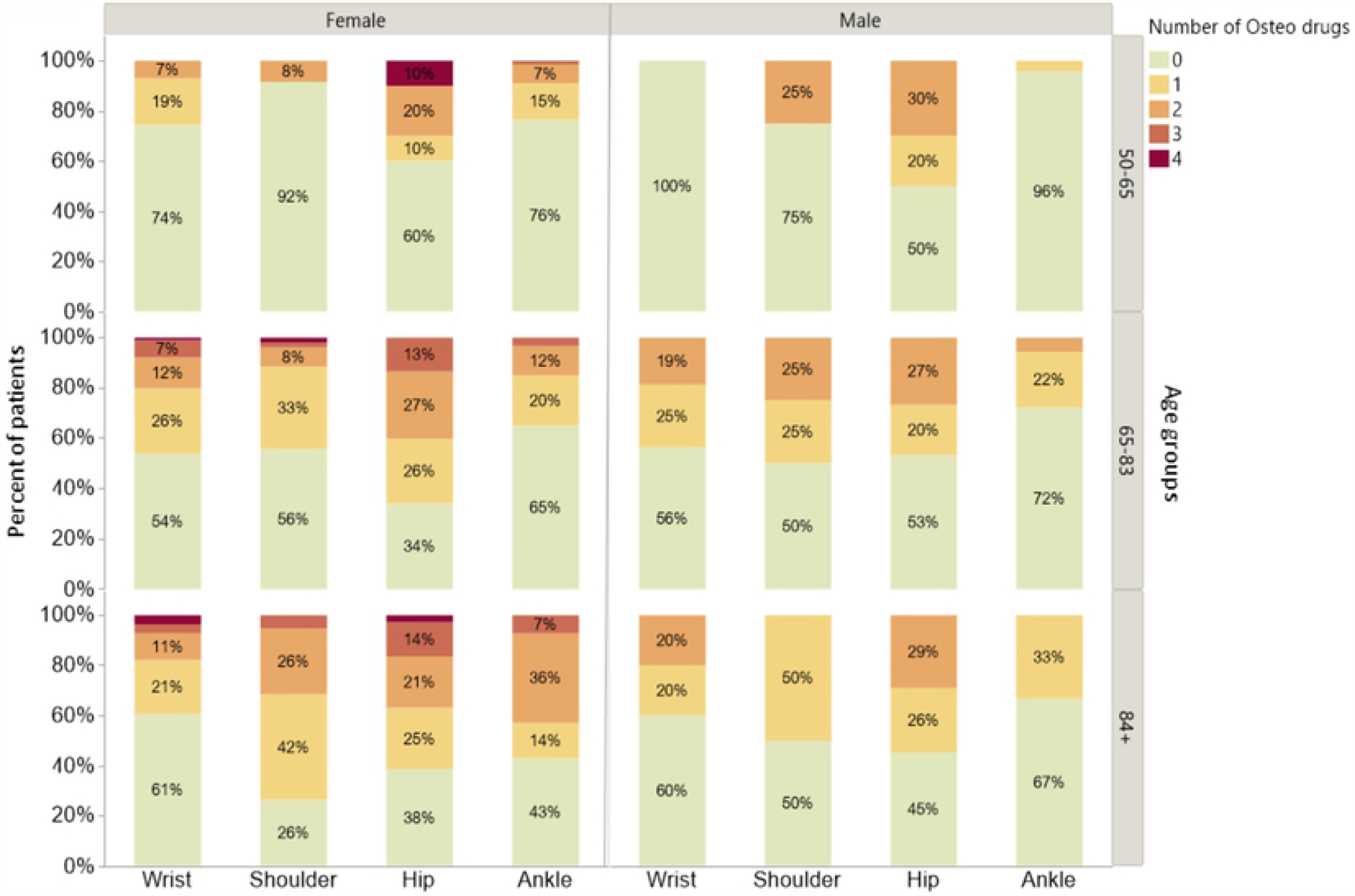
Percent of patients using osteoporosis medications sub-divided by sex and age groups. Zero (0) value in the figure represents patients that were not taking any medications for osteoporosis. It is evident that males were using less number of medications (if any) than females and this was noticeable for hip fractures. For females, the largest increase across age groups in the percent of patients taking medications was for shoulder fractures. Please refer to the main text for odds ratios.

Regardless of fracture site and age, females had a significantly higher odds of using osteoporosis medications than males: 1.77 (95% CI, 1.21 to 2.58). Patients presenting with hip fractures at any age had the highest odds of taking osteoporosis medication compared to patients with fractures in any other body area: 2.66 (95% CI, 1.94 to 3.68). The odds for patients presenting with shoulder fractures to be on osteoporosis medications was very similar to other fractures at 1.13 (95% CI, 0.73 to 1.73). The older the patient, the higher the probability of patients taking osteoporosis medications. The odds for osteoporosis medication use in patients in the 84 years and older group compared to patients in the 50-65 years group was 4.92 (95% CI, 3.12 to 7.82). Similarly, patients in the 66-83 years group had odds of taking medication of 3.75 (95% CI, 2.29 to 5.19) than the younger group.

## Discussion

Contrary to our hypothesis, the use of medications to prevent osteoporosis in patients that present fragility fractures were comparable to previously reported national averages [19]. Within the set of patients taking osteoporosis medications, the use of calcium, vitamin D, and bisphosphonates constituted more than 80% of the consumption. When calcium and vitamin D were not considered, the percent of patients on osteoporosis medications dropped by 25% (from 42 to 16.7%) but remained above the national averages of 10% for years 2010-2011 [19]. Females were more likely to take osteoporosis medications, and this probability improved with increased age. In our population, male patients presenting with fractures and treated with osteoporosis medications were very low. This finding is similar to previous reports which are partially explained due to underdiagnosed and under-treatment of males for osteoporosis [20,21]. To our knowledge, this is the first time that a study identifies the use of medications for osteoporosis within the southernmost region of the United States, with a high prevalence of Hispanic/Latino heritage.

When we examined the socioeconomic status and self-reported race/ethnicity, reports have shown that individuals at an extreme socioeconomic disadvantage are very vulnerable to relatively low bone mineral density [22]. The Rio Grande Valley population is well documented to have low socioeconomic status with a median income of only $39,000 dollars. The Rio Grande Valley is also classified as a medically underserved population with a prevalence of 30% for uninsured/underinsured patients [23]. Despite these well documented socioeconomic conditions, the use of osteoporotic medication was not lower than the national average. However, it is essential to consider that vitamin D and calcium, which can be obtained at a low cost and without prescription, were the predominant medications used in our population. Within our cohort, patients taking medications for osteoporosis were older and had a higher prevalence of comorbidities than patients not taking medications for osteoporosis. The need for closer multidisciplinary medical treatment for patients with increased age and multiple comorbidities produce more medical interventions and closer monitoring, allowing for identification of the patient is at risk of osteoporosis and medical treatment interventions.

A study looking at bone turnover in Mexican Americans who also have type 2 diabetes found lower bone turnover in men with diabetes and poor glycemic control [24]; hence, screening for osteoporosis in Mexican Americans to prevent fractures should be highly considered [25]. Native Americans, White and Hispanic women remain among the highest for fracture risk than other ethnic groups (Black, Asian)[26]. Given that the Rio Grande Valley population is medically underserved and has a high prevalence of type 2 diabetes, we believe that there could be an increased risk of osteoporosis-related fractures in our community. However, this information has never been collected nor reported.

We observed a discrepancy for prior osteoporosis diagnosis within our data with the number of patients taking the medications. This discrepancy might be produced by a lack of proper documentation within the medical records or simply by the retrospective nature of the data. It is possible that patients might have been diagnosed at a primary care facility not associated with our hospital system; thus, documentation of DXA scans or diagnosis is not within our system’s electronic medical record. However, it should be noted that in the mid-nineties, the use of bone mineral density scans for osteoporosis was not recommended [27], but this view has been challenged, and DXA remains the “gold standard” for osteoporosis diagnosis [28,29]. An additional possibility to consider is that the patients initiate the reported use of calcium and vitamin D on their own, as part of a multi-vitamin regime, or only as a physician’s recommendation due to their advanced age. The latter is supported by a previous study indicating that for Hispanics, information regarding medication use and adherence is more readily received from the doctor than from any other source of information: “the doctor is still king” [30]. Future prospective studies should address the process of diagnosis and reporting, both by primary care and specialty doctors.

### Study limitations

Retrospective cohort studies provide a quick estimate when no previous data on the topic exist, especially for specific populations, but it also carries a series of limitations. Information regarding the length that the patients have been taking the medication was not collected, nor the effectiveness of such medications. While the number of patients taking medications in the current study is not suggestive of underreporting, it is always possible that patients are taking calcium plus vitamin D as part of their regular multi-vitamins but do not consider vitamins as “medicine” or “treatment.” Hence the possibility of underreporting is impossible to rule out. An attempt to verify previous fracture history was initiated, but we ruled out acquiring this information given that our region has three large hospitals. The possibility of patients seeking care for a prior fracture at a different facility is high. Thus future studies should consider multi-institutional design, allowing the recording of this type of information. We acknowledge an intrinsic bias in the study design by selecting patients that already present with a fracture.

However, the presence of a fragility fracture is one of the critical factors for the diagnosis of osteoporosis [31]. Based on the new practice guidelines for endocrinologists, the current patient population would fall on the very high-risk group based on the presence of a fracture within the previous 12 months [29]. Despite the limitations, the current study sets the stage for designing prospective interventions in high-risk groups for osteoporotic fractures.

## Conclusions

Osteoporosis continues to be an underdiagnosed and undertreated disease. In our cohort, despite the high use of prophylactic medication among the most elderly patients, the usage of medications in the younger population continues to be minimal. Within the male population, the usage and diagnosis of osteoporosis continue to be almost nonexistent. Despite slightly higher use of prophylactic medication than national standards, the percentage of patients taking medication still falls under desired levels, especially considering that only 16% of the patients took medication once vitamin D plus calcium were removed from the comparison. It is essential to recognize there is still significant work to promote consciousness, improve diagnoses, and encourage early use of prophylactic medications.

## Acknowledgments

We acknowledge the valuable contribution of the following individuals in the process of data extraction and data mining, all prior or current employees of DHR Health: Mr. Miguel de la Rosa, Mr. Jason Martinez, and Mr. William Noonan. We appreciate the valuable comments on the manuscript from Dr. Lisa Trevino and Mr. Peter Roberge from DHR health Institute for Research and Development.

## Ethics approval

All procedures performed in studies involving human participants were in accordance with the ethical standards of the institutional and/or national research committee and with the 1964 Helsinki declaration and its later amendments or comparable ethical standards.

## Informed Consent and Approval

This retrospective study received approval on February 3, 2020, by the DHR Health Institute for Research and Development IRB under the expedited mechanism of review. A full waiver of consent to participate was submitted and approved by the IRB, specifying the retrospective nature of the study and the age of the participants to be included. All data was collected and analyzed de-identified.

## Data Availability

The patient information data used to support the findings of this study are restricted by the DHR Health Institute for Research and Development Institutional Review Board to protect patient privacy and comply with HIPPA laws. De-identified data sets are available from the corresponding author, Dr. M. Serra-Torres, for researchers who meet the criteria for access to confidential data. Dr. Serra can be reached by email at.

## Author’s contributions

MJST conceived the original idea, designed the protocol, and contributed to the writing of the first draft. ATR, analyzed the data, prepared figures and the first draft of the manuscript. Both authors discussed the results and conclusions and approved the final version of the submitted manuscript.

